# Comprehensive functional profiling of long non-coding RNAs through a novel pan-cancer integration approach and modular analysis of their protein-coding gene association networks

**DOI:** 10.1101/254722

**Authors:** Kevin Walters, Radmir Sarsenov, Wen Siong Too, Roseanna K. Hare, Ian C. Paterson, Daniel W. Lambert, Stephen Brown, James R. Bradford

## Abstract

Long non-coding RNAs (lncRNAs) are emerging as crucial regulators of cellular processes in diseases such as cancer, although the functions of most remain poorly understood. To address this, here we apply a novel strategy to integrate gene expression profiles across 32 cancer types, and cluster human lncRNAs based on their pan-cancer protein-coding gene associations. By doing so, we derive 16 lncRNA modules whose unique properties allow simultaneous inference of function, disease specificity and regulation for over 800 lncRNAs. Remarkably, modules could be grouped into just four functional themes: transcription regulation, immunological, extracellular, and neurological, with module generation frequently driven by lncRNA tissue specificity. Notably, three modules associated with the extracellular matrix represented potential networks of lncRNAs regulating key events in tumour progression. These included a tumour-specific signature of 33 lncRNAs that may play a role in inducing epithelialmesenchymal transition through modulation of TGFβ signalling, and two stromal-specific modules comprising 26 lncRNAs linked to a tumour suppressive microenvironment, and 12 lncRNAs related to cancer-associated fibroblasts. At least one member of the 12-lncRNA signature was experimentally supported by siRNA knockdown, which resulted in attenuated differentiation of quiescent fibroblasts to a cancer-associated phenotype. Overall, the study provides a unique pan-cancer perspective on the lncRNA functional landscape, acting as a global source of novel hypotheses on lncRNA contribution to tumour progression.

**Author Summary:** The established view of protein production is that genomic DNA is transcribed into RNA, which is then translated into protein. Proteins play a critical role in shaping the function of each individual cell in the human body yet they represent less than 2% of human genomic sequence whilst up to 90% of the genome is transcribed. To explain this disparity, the existence of thousands of long non-coding RNAs (lncRNAs) has emerged that do not encode proteins but perform function as an RNA molecule. Most lncRNAs have yet to be assigned a specific biological role, so to address this we apply a novel computational approach to characterise the function of >800 lncRNAs through consistent association with protein coding genes across multiple cancer types. By doing so, we discover 16 “modules” of closely related lncRNAs that share broad functional themes, the most compelling of which consists of 12 lncRNAs that could regulate activation of specific cells neighbouring the tumour, leading to accelerated tumour progression and invasion. Overall, the study provides the most robust view of the lncRNA-protein coding gene landscape to date, adding to growing evidence that lncRNAs are key regulators of cancer, and have therapeutic potential comparable to proteins.

## Background

The advent of high-throughput genomic technologies such as Next Generation Sequencing (NGS) has led to remarkable progress over the last decade in detecting novel transcripts, many of which have no apparent protein-coding capacity. A significant proportion of these non-coding species are long non-coding RNAs (lncRNAs), which typically exceed 200 nucleotides in length, and function through a variety of mechanisms including remodelling of chromatin, transcriptional co-activation/repression, protein inhibition, post-transcriptional modification, or decoy. They are now emerging as crucial regulators of cellular processes and diseases, and their aberrant transcription can lead to altered expression of several important target genes involved in cancer [1], resulting in tumour progression and poor prognosis [2][3][4][5][6].

Despite advances, the vast majority of lncRNAs identified through large-scale efforts such as GENCODE [7] and MiTranscriptome [8] remain poorly understood. To address this gap, several computational approaches have been developed with the ability to assign putative function to thousands of lncRNAs simultaneously by exploiting the widespread availability of cancer genomic data [9][10]. These methods typically employ a “guilt-by-association” strategy, deriving a prediction based on a common expression pattern between the lncRNA and a biological process or pathway [11]. More recent efforts attempt to strengthen predictions by combining transcriptomic data across multiple cancer [12][13][14][15], or normal tissue types [16]. However, whilst representing important advances, these have so far employed limited integration strategies, either seeking consensus across separate lncRNA signatures derived from a small number of cancer types [12][13], correlation across a single dataset against a restricted set of cancer genes [15], or focusing on a natural antisense transcripts only [16].

To address these shortcomings, we have developed a unique workflow to integrate expression associations between lncRNA and protein coding (PC) genes across 32 different cancer types from The Cancer Genome Atlas (TCGA) to provide a more robust lncRNA-PC association network than can be derived from any single cancer type alone. The workflow incorporates three novel aspects: (1) An Expectation Maximisation (EM) algorithm for estimating the correlation between a lncRNA and PC gene that specifically addresses low lncRNA expression relative to PC gene expression. (2) A statistical method for integrating lncRNA-PC correlations across multiple cancer types to derive a single multicancer association (MCA) score between each lncRNA and PC gene, allowing subsequent construction of a single pan-cancer lncRNA-PC gene network. (3) A unique application of Weighted Gene Correlation Network Analysis (WGCNA) [17] to the lncRNA-PC MCA score network allowing its de-convolution into lncRNAs that share consistently similar expression profiles across multiple cancers, henceforth termed “modules”.

Through detailed characterisation of these modules, we provide the most comprehensive pan-cancer assessment of lncRNA-PC gene expression associations to date, allowing simultaneous hypothesis generation on lncRNA function, disease specificity, and transcription factor regulation. More specifically, the unique global perspective of our modular approach reveals the potential for both coordinated and antagonistic lncRNA expression to underpin disease pathway regulation, and new insights into the role of lncRNAs in the tumour microenvironment.

## Results and Discussion

### A workflow to identify lncRNA modules based on their pan-cancer protein coding gene associations

The workflow is divided into two main stages (Figure 1). In the first stage, RNA-Seq expression estimates of each lncRNA and PC gene annotated by GENCODE [7] were inspected across all 32 cancer types (Table S1), and those that failed to achieve sufficient expression signal in any cancer type were removed (see *Methods* for specific criteria). 1833 lncRNAs expressed in at least one cancer type remained after filtering. An EM algorithm was then applied to estimate a pan-cancer correlation coefficient 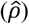 between the expression profiles of each of these 1833 lncRNAs and 17088 PC genes across 1≤n≤32 cancer types in which the lncRNA expression threshold had been met. The approach was specifically developed to handle instances where lncRNA expression is either low, absent, or undetectable across an excessive number of samples, even if initial expression level criteria had been met. Bootstrapping then quantified the uncertainty of 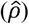 in the form of a standard error 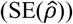, from which a multi-cancer association (MCA) score was derived: 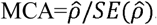. The MCA score calculation was repeated across all PC genes to generate an MCA profile of 17088 scores for each of the 1833 lncRNAs. The lncRNA-PC gene combination achieving the highest MCA score represented the strongest pan-cancer expression association for that lncRNA. Collectively the profiles formed a matrix of 1833x17088 MCA scores. Full details of the derivation of 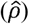 and 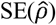 are described in *Methods*.

**Fig 1:**
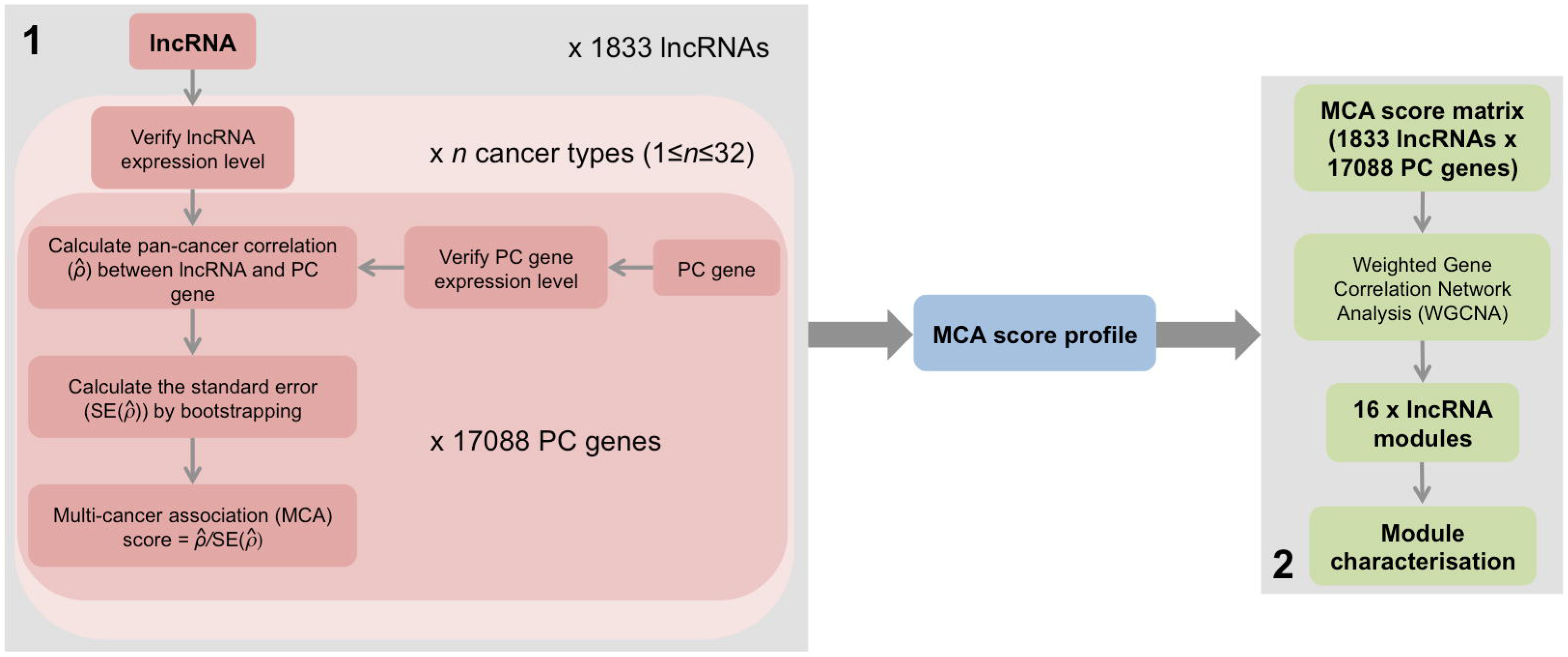
Schematic of workflow to identify lncRNA modules based on their pan-cancer PC gene associations. (1) Derivation of a multi-cancer association (MCA) score profile for 1833 lncRNAs that pass expression level criteria, and (2) identification of 16 modules of highly correlated lncRNAs using Weighted Gene Correlation Analysis (WGCNA) and a matrix of 1833 lncRNA MCA score profiles across 17088 PC genes as input.

In stage two, we applied WGCNA [17] to the MCA score matrix. WGCNA is often used as a dimensionally reduction method in genomics, typically applied to gene expression networks of several thousand genes to identify a small number of modules of related genes whose expression profiles are highly correlated. Each module is represented by an eigen-gene, which can be used to correlate modules with meta-data such as clinical traits. The correlation of a gene’s expression profile with a module eigengene (ME) provides a measure of significance of the relationship between gene and module. Here, we adapted WGCNA to generate “eigen-lncs”, which are analogous to eigen-genes, to identify 16 modules of lncRNAs with highly correlated MCA score profiles (Figure 2A; Table S2). An important advantage of this approach is that the eigen-lnc coefficients attributed to each PC gene (henceforth referred to PC-module association or PC-MA values) can be used as a surrogate for the strength of the relationship between PC gene and eigen-lnc (Table S3). This allowed for functional traits representative of each module to be identified since each module is related to a set of highly annotated PC genes. Here, we defined PC genes achieving PC-MA>0.02 as “pro-module” (PC genes whose mRNA expression is consistently positively correlated with members of the module), and PC-MA<-0.02 as “anti-module” (PC genes consistently negatively correlated with the module).

**Fig 2:**
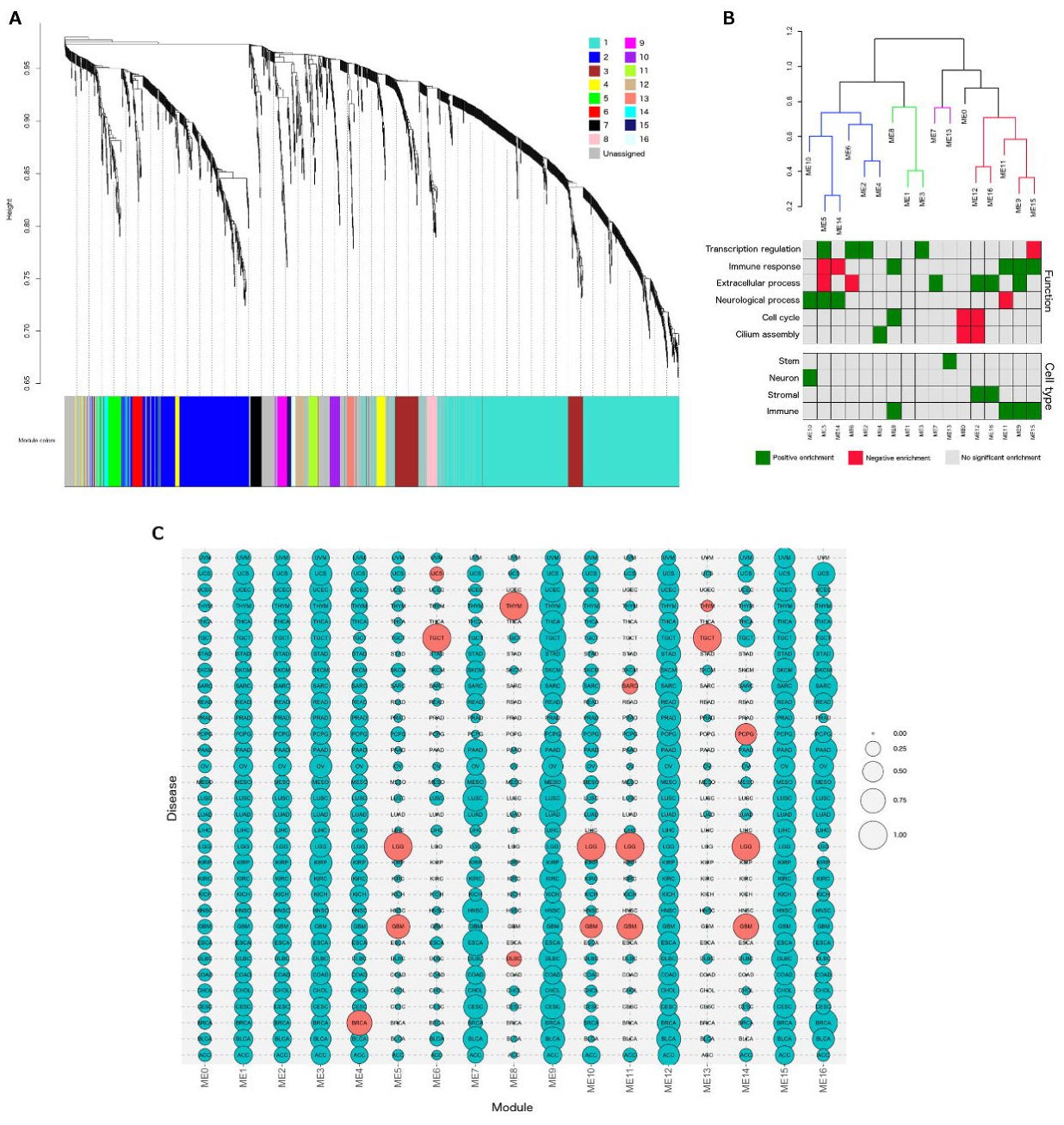
Module characterisation. A. Dendrogram showing hierarchical clustering of lncRNAs based on MCA score profile. Branches of the dendrogram correspond to modules, with lncRNAs in each module assigned the same colour (indicated by the colour band below the dendrogram). LncRNAs not assigned to a module are coloured grey. B. Clustering dendrograms of module eigen-lncs. Meta-modules are defined at height cut-off of 0.80 and indicated by different colours. Below the dendrogram, functional and cell type signatures of each module are indicated, with green corresponding to significant positive enrichment and red to significant negative enrichment. C. Bubble chart showing cancer type specificity of each module. Size of bubble indicates the proportion of module-associated lncRNAs that meet the expression detection threshold in each cancer type. Red bubbles indicate outlier cancer types (>1.5 times the interquartile range above the upper quartile). A description of the cancer type codes is given in Table S1.

### Module characterisation

#### Common functional traits

Functional enrichment analysis of pro-module PC genes revealed striking properties of lncRNA-PC gene associations (Table 1, Figure 2B, Table S4). Primarily, modules could be grouped into four functional signatures: immune, extracellular, transcription regulation, and neurological, broadly corresponding to four sets of positively correlated eigen-lncs or “meta-modules” (Figure 2B, Figure S1). Only ME4 (cilium assembly; *p*=9.38E-08) and ME13 (stem cell signature; *p*=4.05E-41) fell outside the general classification. ME8 was enriched for both cell cycle (*p*=7.49E-30) and immune-associated genes (*p*=1.54E-29). Four of the top six largest modules ME2, ME3, ME5 and ME6 comprising the majority of lncRNAs (524/822) were associated with transcriptional regulation (Table 1), possibly reflecting the common role of lncRNAs in chromatin structure modification and control of PC gene expression [18]. The smaller modules were typically related to more specific signatures, including four modules associated with the immune system (ME8, ME9, ME11, ME15), three with the extracellular matrix (ME7, ME12, ME16), and three with neurological processes (ME5, ME10, ME14). No coherent functional signature could be established for the largest module (ME1) of 723 lncRNAs, and 288 lncRNAs were allocated to a pseudo-module (ME0) since their module membership could not be established. Overall, a putative pan-cancer functional association could be assigned to 822 lncRNAs by our approach.

**Table 1:** Module features

| Module | Number of lncRNAs | Functional signature |
| --- | --- | --- |
| ME0 | 288 | None |
| ME1 | 723 | None |
| ME2 | 317 | Transcriptional regulation |
| ME3 | 128 | Transcriptional regulation |
| ME4 | 56 | Cilium assembly |
| ME5 | 46 | Transcriptional regulation / Neurological |
| ME6 | 33 | Transcriptional regulation |
| ME7 | 33 | Extracellular |
| ME8 | 32 | Immune / cell cycle |
| ME9 | 31 | Immune / extracellular |
| ME10 | 30 | Neurological |
| ME11 | 29 | Immune |
| ME12 | 26 | Extracellular |
| ME13 | 21 | Stem cell |
| ME14 | 16 | Neurological |
| ME15 | 12 | Immune |
| ME16 | 12 | Extracellular |

#### Tissue type specificity

The functional themes of several modules reflected the cancer or normal tissue type specificity of their lncRNAs (Figure 2C, Table S5) [19]. As expected, neurological-associated ME5, ME10, and ME14 were highly specific to brain cancers, and all 21 lncRNAs of stem cell associated ME13 were detected in testicular germ cell tumours (TGCT), consistent with the notion that TGCT cells are derived from normal germ cells with distinct stem cell characteristics [20]. ME6 was also highly specific to TGCT, and whilst there was no significant association with a stem cell signature, it included the lncRNA *LINC-ROR*, which modulates reprogramming of fibroblasts to a pluripotent stem cell state [21]. Likewise, the enrichment of ME8 for immune processes such as lymphocyte activation (*p*=1.54E-29) reflected its specificity for thymoma, and the origins of this cancer type in the thymus gland. Interestingly, ME8 was also associated with the cell cycle (*p*=7.49E-30), which is emerging as a potential prognostic indicator in thymoma [22]. No disease bias was observed in transcriptional regulation-associated modules ME2 and ME3, immune-associated ME9 and ME15, and extracellular matrix-associated ME7, ME12 and ME16, suggesting that lncRNAs in these modules contribute to fundamental cellular processes common to most cancer types.

### Detailed characterisation of the extracellular-associated modules

Given their pan-cancer expression, and current poor understanding of the role of lncRNAs in extracellular processes, we were keen to dissect modules ME7, ME12 and ME16 further, and generate hypotheses on their potential function in supporting tumour progression.

#### FOS/JUND transcription factor binding site enrichment in ME7

To establish whether lncRNAs in each of the extracellular modules share a common promoter, we performed a *de novo* search for sequence motifs in regions 1000bp upstream of the lncRNA transcription start site (TSS). Whilst there was no evidence for transcription factor binding enrichment in ME12 and ME16, a top scoring motif achieving >95% similarity with FOS and JUND transcription factor binding sites [23] was observed in 18/33 lncRNAs of ME7, (Figure 3A; Tables S6a-d). There was no evidence for enrichment of the FOS/JUND motif in the other 15 modules.

**Fig 3:**
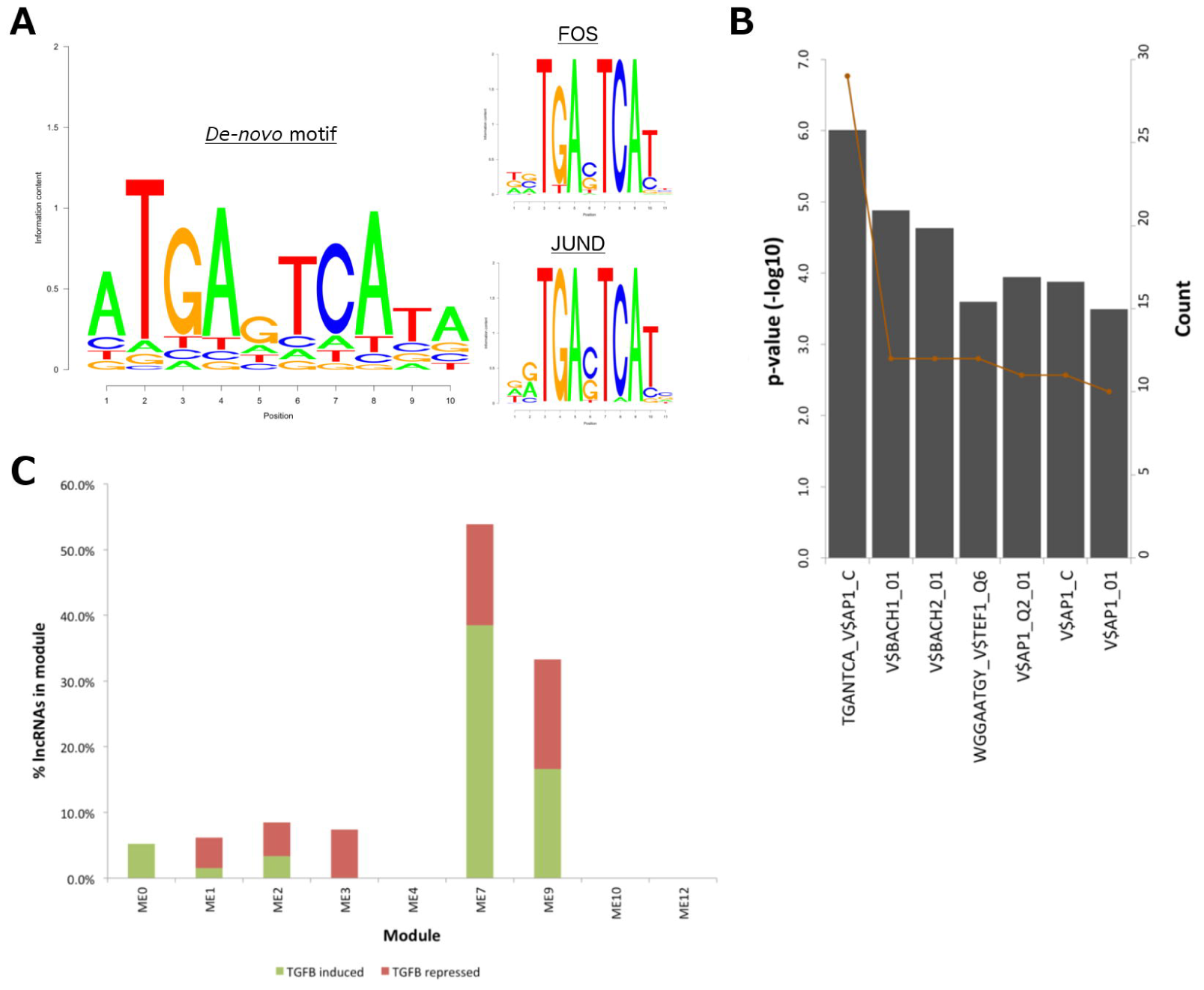
Evidence of c-Fos/c-Jun regulation in ME7. A. Top-scoring *de-novo* motif in the region 1000bp upstream of lncRNA transcription start site, and top two most similar JASPAR [29] transcription factor motifs. B. Enrichment of AP1-like binding sites in pro-module PC genes of ME7 according. C. Proportion of lncRNAs in each module regulated by TGF‐β and occupied by SMAD3 according to [32].

Underpinning this discovery, pro-module PC genes of ME7 were enriched for the binding site of activator protein-1 (AP-1) (*p*=9.80E-07; Figure 3B; Table S7), a transcription factor dimer of Jun and the Fos family of basic leucine zipper domain proteins, with FOS-like antigen 1 (*FOSL1*) achieving the highest PC-MA value of 0.036 (Table S3). Moreover, strong binding of both c-Jun and c-Fos to the promoter region of ME7 lncRNA, *RP11-554I8.2* (also known as *LINP1*), has recently been confirmed in triple negative breast cancer cell lines [24].

Since c-Jun and c-Fos are known to co-operate with mothers against decapentaplegic homolog (SMAD) proteins to mediate transforming growth factor beta (TGFβ) signalling at AP-1 binding sites [25], we compared ME7 with two studies on SMAD targets [26][27]. Firstly, overlap with [26] revealed 39% (13/33) of ME7 lncRNAs are expressed in human hepatic stellate cells (HSC) (Table S8), representing the highest enrichment compared to the other modules. Of these, 53% (7/13) are potential targets for SMAD3 representing significant enrichment (*p*=0.01 by hyper-geometric test), and either induced (39%; 5/13) or repressed (15%; 2/13) by TGFβ signalling (Figure 3C, Table S8). Similarly, comparison with [27] showed that the promoters of 13 of the top 20 ME7 pro-module genes could be occupied by either SMAD2 or SMAD3.

TGFβ induces epithelial-mesenchymal transition (EMT) in tumours via activation of SMAD proteins [28], which translocate into the nucleus and regulate transcription of TGFβ target genes [29]. Furthermore, since SMADs have low affinity for DNA, it is crucial they interact with cofactors such as AP-1 [30] to achieve target specificity. Exploring a potential link between ME7 and EMT induction via TGFβ signalling, we observed significant enrichment (*p*=8.74E-20) for an EMT signature in pro-ME7 PC genes that included *SNAI2* (PC-MA=0.025) and *TGFβ1* (PC-MA=0.021). Pro-ME7 PC genes also included *HMGA2* (PC-MA=0.024), a downstream effector of TGFβ during EMT [31], and *FOSL1*, whose protein product Fos-related antigen 1 (Fra-1) is implicated in EMT through modulation of TGFβ expression [32]. Taken together, our results indicate that lncRNAs of ME7 play a role in the induction of EMT via convergence of AP-1 and SMAD proteins at their promoters and regulation of TGFβ signalling.

#### Determination of the tumour stromal specificity of ME12 and ME16

We noted that ME12 and ME16 shared a number of pro-module PC genes (Figure 4A) and achieved significant correlation between their eigen-lncs (*r*=0.57). In addition, both pro-module PC gene sets of ME12 and ME16 overlapped significantly (*p*<0.05 by hyper-geometric test) with a stromal cell signature [33], incorporating 24% (32/136) and 35% (48/136) signature genes respectively. By contrast, no overlap was observed with ME7.

**Fig 4:**
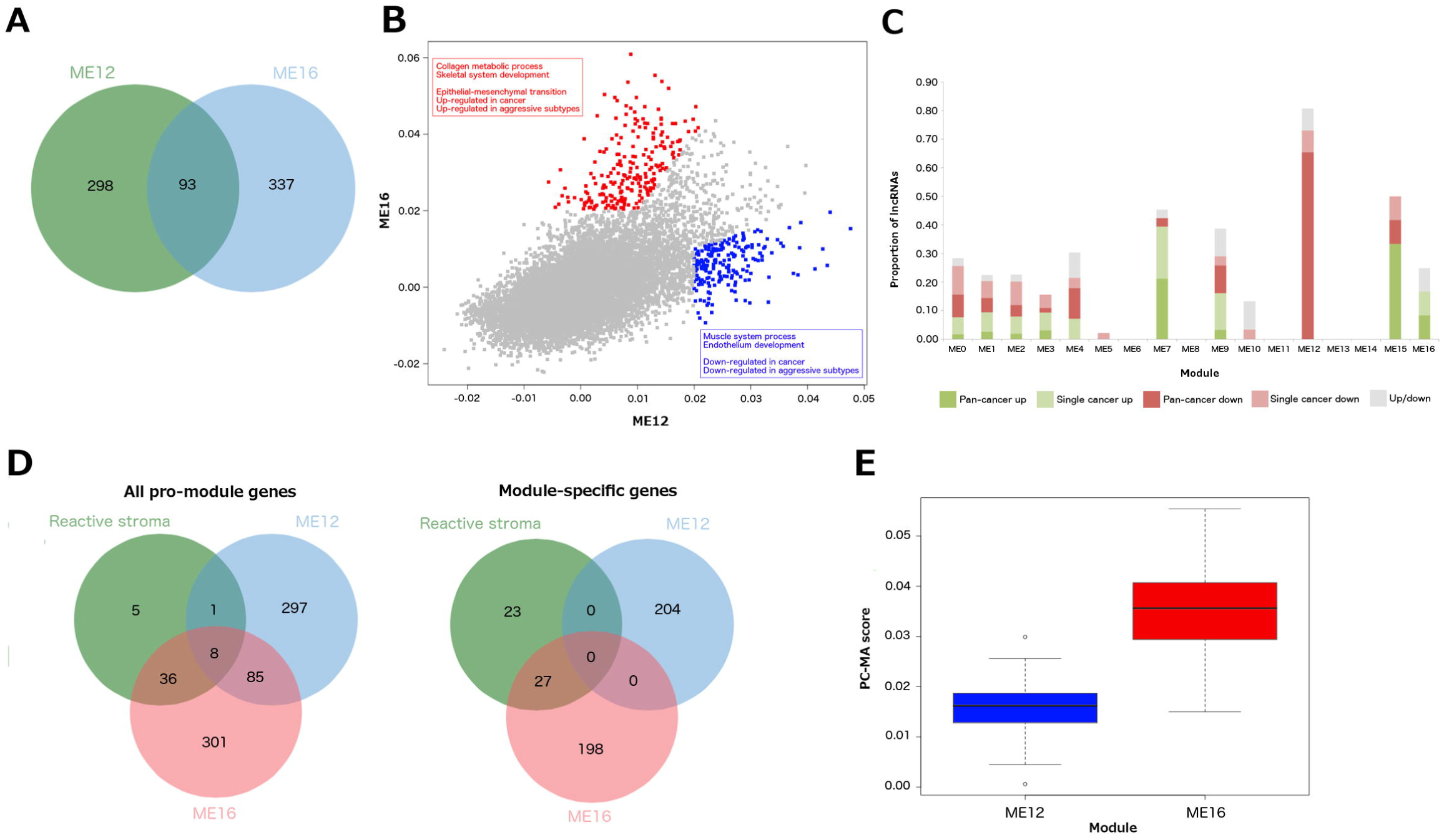
Differentiation of the extracellular-associated modules ME12 and ME16. A. Venn diagram to show overlap of pro-module PC genes between ME12 and ME16. B. Scatterplot of ME16 versus ME12 PC-MA values. ME16-specific PC genes are indicated in red, ME12 in blue with corresponding text highlighting signature enrichments in module-specific lists. C. Proportion of lncRNAs potentially dysregulated in cancer within each module. D. Venn diagrams showing overlap of all pro-module genes and module specific genes with a reactive stroma signature [48]. E. Boxplot comparing PC-MA distribution across reactive stroma signature genes between ME12 and ME16.

We explored the potential stromal specificity of ME12 and ME16 further by using a novel approach to generate a putative list of 300 stromal cell specific (SCS) lncRNAs frequently detected in stromalcontaining clinical samples but not in pre-clinical models that consist almost exclusively of tumour cells (Table S9a; see *Methods*). Both ME12 and ME16 contained an abundance of SCS lncRNAs, achieving 60% (15/25) and 53.6% (7/11) overlap respectively (Table S9b). By contrast, only 1/27 (3.7%) lncRNAs in ME7 were classed as SCS. These results provide strong *in silico* evidence that expression of lncRNAs in ME12 and ME16 is specific to the tumour stroma.

#### Functional dissection of stromal-specific modules ME12 and ME16

To define more precise roles for the lncRNAs of ME12 and ME16, we identified 204 “ME12 specific” and 226 “ME16-specific” PC genes that achieved a PC-MA fold difference>2.00 with the corresponding PC gene in ME16 and ME12 respectively (Figure 4B; Table S10a and S10b).

Comparison with signatures from MSigDB [34] revealed ME12-specific PC genes were consistently down-regulated in cancers including prostate (*p*=1.79E-34) and colorectal (*p*=3.10E-18), and advanced disease such as metastatic prostate cancer (*p*=1.07E-12). They were also down-regulated in aggressive cancer subtypes such as luminal-B (*p*=2.38E-17) and basal-like breast cancers (*p*=9.12E-09). In contrast, ME16-specific PC genes were consistently associated with hallmarks of tumour progression such as EMT (*p*=3.90E-67), and up-regulated in aggressive subtypes such as basal-like breast cancer (*p*=1.00E-11).

Further evidence for under-expression of ME12 lncRNAs in cancer was provided by a systematic comparison of lncRNA expression between tumour and normal samples across 14 cancers types (see *Methods*). Each lncRNA was classified as “pan-cancer up” or “pan-cancer down” if differential expression was consistent across more than one cancer type, “single cancer up” or “single cancer down” if observed in a single cancer type, or “both” if the lncRNA was differentially expressed in both directions across different cancer types (Table S11a and 11b). 73% (19/26) of lncRNAs in ME12 were classed as “pan-cancer down” or “single cancer down” (Figure 4C, Table S11c). These included the known tumour suppressor maternally expressed gene 3 (*MEG3*) [35], which was under-expressed in 4/14 cancers represented in our dataset. By contrast, only three lncRNAs in ME16 were classed as differentially expressed, and none as either “pan-cancer down” or “single cancer down”.

Interestingly, 39% (13/33) of lncRNAs in the third extracellular-associated module ME7 were classed as either “pan-cancer up” or “single cancer up”, with >70% of these over-expressed in head and neck squamous cell carcinoma (HNSCC). This was concurrent with the strong association between ME7 and *FOSL1*, which is consistently over-expressed in HNSCC [36].

#### Comparison of ME12 and ME16 with a reactive stroma signature

The above findings led us to compare ME12 and ME16 with a reactive stroma signature [37]. 44/50 genes in the signature overlapped with pro-module PC genes of ME16 compared to only 9/50 genes with ME12 (Figure 4D). These included fibroblast-activation protein (*FAP*; PC-MA=0.06), an established cancer-associated fibroblast (CAF) marker, periostin (*POSTN*; PC-MA=0.05) a gene implicated in metastasis [38], and members of the collagen family such as *COL5A2* (PC-MA=0.05), *COL6A3* (PCMA=0.05), *COL10A1* (PC-MA=0.04) and *COL6A1* (PC-MA=0.04). Considering only module-specific genes, 27/50 genes in the signature overlapped with ME16 but none with ME12 (Figure 4D). Moreover, signature genes achieved significantly higher PC-MA values with ME16 than ME12 (*p*=1.59E-20 by Student’s *t*-test; Figure 4E, Table S12). Taken together, our results strongly suggest that ME16 lncRNAs are markers of an activated stromal phenotype that promotes tumour progression, whereas ME12 lncRNA expression supports a tumour suppressive microenvironment.

#### esiRNA knockdown of ME16 lncRNAs

Given the strong evidence for their stromal cell specificity, and association with activated stroma, we took forward two lncRNAs of ME16 (*AC093850.2* and *RP11-626H12.2*) to experimentally assess their role in the tumour microenvironment, alongside a lncRNA not associated with this module (*RP1-122P22.2* from ME2) and a non-targeting esiRNA (Evf-2) as negative controls. The nearest upstream neighbour of *AC093850.2* is fibronectin (*FN1*), thus providing a potential example of a *cis*-relationship between a known fibroblast marker and lncRNA. To our knowledge, there is no evidence that the protein coding neighbours of *RP11-626H12.2* play a direct role in CAF activation.

We used an established experimental model of CAF differentiation [39] that uses TGF-β1 to activate human primary fibroblasts, assessed as induction of alpha-smooth muscle actin (αSMA) a commonly used CAF marker (Figure 5A). CAFs are the major cell type in the tumour microenvironment and are known to play a role in the invasion and metastasis of tumour cells [40][41]. There is strong evidence showing association between CAFs and poor prognosis in several types of cancers [42]. Figure 5A shows that the knock-down of AC093830.2 has an effect on cell number, but is not completely required for cell viability. We observed that TGF-β1-mediated activation of fibroblasts (as assessed by the number of cells harbouring αSMA-positive stress fibres determined by immunofluorescent labelling and high content microscopy) is impaired when expression of both candidate lncRNAs, but not lncRNA from different functional modules, is reduced in human fibroblasts using specific esiRNAs (Figure 5B). This reduction in TGF-β1-mediated stress-fibre formation reached statistical significance for one of the lncRNAs, *AC093830.2*, when compared to the response in the presence of esiRNA targeting a gene not expressed in these cells (Evf-2).

**Fig 5:**
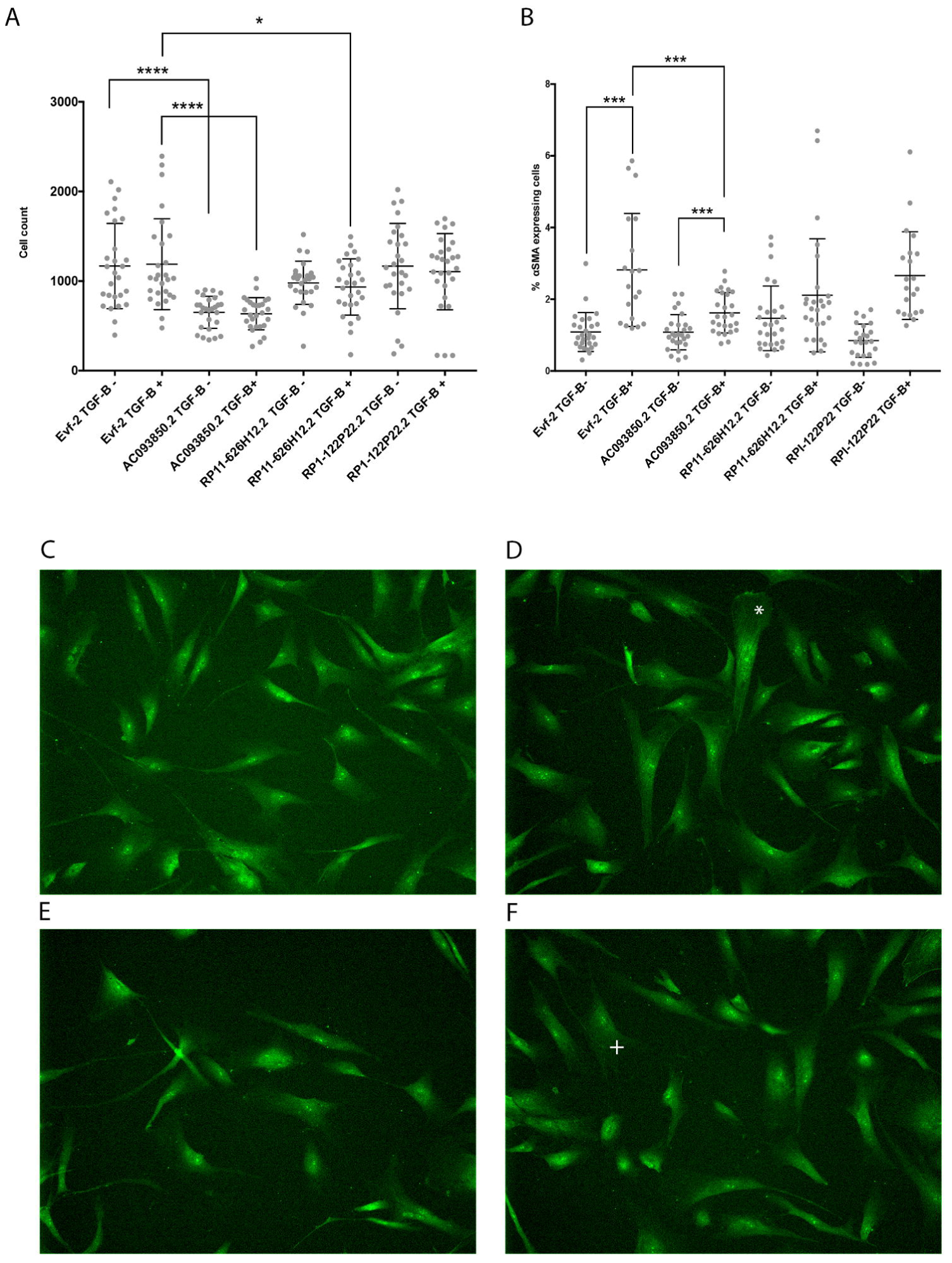
A candidate lncRNA is able to attenuate TGF-β1 1-induced fibroblast differentiation. Human primary fibroblasts were transfected with esiRNA targeting two candidate lncRNAs (*AC093850.2* and *RP11-626H12.2*), a transcript not expressed in human cells (Evf-2) and a lncRNA from a different module not predicted to influence fibroblast differentiation (*RP1-122P22.2*). Each experiment consisted of nine technical and three biological replicates. A. Cells were dispensed into 384 well plates, reverse transfected with esiRNAs, incubated for 2 days knock-down and then stimulated or not with TGF for 24 hours. Images were acquired using a MetaXpress Micro x2 objective and cells identified using the nuclei stain Hoechst, and segmented using MetaXpress software. Data processed in Excel and Prism7. B. The protocol was identical to that of A, but the cell were stained with SMA antibody after fixation, and imaged using the x20 objective. Positive CAFs were identified on the formation of *de novo* SMA-positive stress fibres and morphological changes using MetaXpress Custom Module Editor. \*\**p*<0.05, \*\*\**p*<0.01, \*\*\*\**p*<0.001. C. Microscope images of unstimulated, control Evf-2 knock-down cells. D. TGF-β1- stimulated, control Evf-2 knock-down cells. The white * indicates a transformed CAF with both morphological and SMA positive fibres. E. Unstimulated, *AC093850.2* knock-down cells. F. TGF-β1-stimulated, *AC093850.2* knockdow5n cells. The white + indicates a cell counted as a CAF with a partial transformation.

In comparison to control (Figures 5C and 5D) and unstimulated *AC093850.2* knock-down (Figure 5E), images of TGF-β1 activated fibroblasts knocked-down with *AC093830.2* RNAi show that the cells are morphologically different to activated wild-type CAFs, and a few cells still show some transformation into the TGF-β1 induced phenotype (Figure 5F, white cross). These results suggest that *AC093850.2* is functionally linked with the differentiation of fibroblasts to a CAF phenotype, and may act in a redundant manner.

Further support for the association of ME16 with the CAF phenotype was provided by analysis of gene expression data from a separate study using the same method of induction of a CAF phenotype [43]. A comparison showed that 20/64 genes over-expressed (log_2_FC>1.50, *p*<1.00E-04) in response to TGF-β1 treatment of HFFF2 fibroblasts are also members of the pro-module gene set of ME16, representing significant overlap (*p*=1.47E-14 by hypergeometric test).

## Conclusion

In this study, we present the most comprehensive examination of the pan-cancer lncRNA expression landscape to date. A key contribution is the development of a novel approach to integrate transcriptome data across multiple cancers, allowing us to generate lncRNA-PC networks and de-convolute lncRNAs into a small number of functionally coherent modules. By doing so, we provide some important insights and hypotheses into the role of lncRNAs in cancer. Principally, lncRNAs can be grouped into just four functional themes based on their associations with PC genes: immune, extracellular, transcription regulation, and neurological.

Whilst a number of modules are clearly driven by the tissue specificity of their lncRNAs, several pan-cancer modules are identified, of which three may represent distinct lncRNA networks associated with extracellular processes that regulate key events in tumour progression. Two of these modules are stromal specific, corresponding to a 26-lncRNA signature associated with a tumour suppressive microenvironment, and 12 lncRNAs with a potential role in cancer fibroblast activation leading to poor prognosis. The third module consists of a tumour-derived signature of 33 lncRNAs that may play a role in inducing EMT through modulation of TGFβ signalling. Adding confidence to our approach, the potential functional regulatory roles of two members of the putative lncRNA CAF signature were validated by experimental modulation in fibroblasts. Interestingly, whilst reduction in TGF-β1-mediated stress-fibre formation was observed for both lncRNAs, it reached statistical significance only for *AC093850.2* (also known as *LINC01614*). The nearest upstream neighbour of *AC093850.2* is fibronectin, a key component of CAF-derived ECM known to influence matrix remodelling associated with metastasis [44]. Therefore, our findings could indicate a lncRNA-mediated control mechanism of fibroblast differentiation via *cis*-regulation of fibronectin by *AC093850.2*.

Since reference to modules alone may mask subtle functional differences that exist between lncRNAs, we encourage researchers to explore the individual lncRNA PC-MA profiles provided as supplementary data (https://figshare.com/s/753cc0df15197b0b9572). Together with the modules, they provide a unique, global compendium from which to generate novel hypotheses and motivate detailed functional studies on lncRNA roles in cancer.

## Methods

### TCGA RNA-Seq data processing

Raw FASTQ sequence files for each solid tumour represented in TCGA were downloaded from the Cancer Genomics Hub (CGHub; https://cghub.ucsc.edu), and reads aligned to the human (GRCh38) genome using StarAlign [45] with no more than three mismatches and only uniquely mapped reads allowed. Reads whose ratio of mismatches to mapped length was greater than 0.10 were also discarded. All other parameters were set to their defaults for unstranded alignment. To reduce possible biases introduced by variable total read counts between samples, tumours achieving <20,000,000 mapped reads were removed. The expression level, based on Fragments Per Kilobase per Million fragments mapped (FPKM), of each gene present in the human (GRCh38) GENCODEv22 annotation file was estimated using Cufflinks with library type defined as “fr-unstranded” and all other parameters set to defaults [46].

Expression values were then batch normalized using COMBAT [47] where appropriate. Only genes annotated as “lincRNA” or “protein_coding” were considered. LncRNAs overlapping PC genes were ignored, as well as genes whose largest transcript is less that 400bp due to potential over-estimation of expression across transcripts less than the average fragment length. We also removed lncRNA and PC genes that failed to achieve sufficient expression signal (mean FPKM + standard deviation (SD))>1.00) across at least one cancer type. The resulting gene-by-sample matrix consisted 17088 PC genes and 2098 lncRNAs. Note that a poly-A selection protocol was used for TCGA RNA-Seq, and so lncRNAs are restricted to these species. Sequencing data for all TCGA cancer types used in this study were processed using the same procedure. The number of tumours across each cancer type is given in Table S1.

### Pan-cancer estimation of the correlation between each lncRNA and PC gene

Visual inspection of the data indicates that a three-component mixture distribution is an appropriate representation. The first two densities can be seen to decay exponentially away from the x and y axes and the third distribution looks bivariate Gaussian (Figure S2). We use the expectation maximisation (EM) algorithm to estimate the parameters of our statistical mixture model. Since we are specifically interested in the correlation coefficient of the bivariate Gaussian density, we estimate the separate parameters of the bivariate Gaussian covariance matrix rather than the whole covariance matrix itself. To exploit the convenience of using sufficient statistics for the parameters, we ensure that the mixture density is in the exponential family. Data across 32 cancer types (indexed by *c*) is used in the maximum likelihood estimation. The three-component mixture density likelihood over the 32 cancer types is:

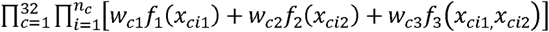

where is the weight for component *j* in cancer type c (such that 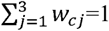 for all cancer types), *n*_*c*_ is the number of samples in cancer type *c*, *x*_*ci*1_ PC gene expression value in cancer type *c.* The three mixture components are

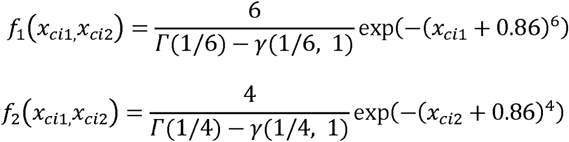

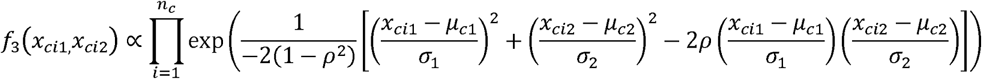

where Γ is the standard gamma function and the lower incomplete gamma function. In order to fit this into the exponential family we assume that the lncRNA and PC gene expression variances for each of the cancer types are identical and defined as 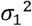 and 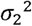. The lncRNA and PC gene expression expectations (μ_c1_ and μ_c2_) are however allowed to vary for each of the cancer types. The correlation coefficient ρ is the parameter of interest.

We use the EM algorithm with updates derived by equating expectations in the usual way. Let 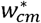 represent the current value of the parameter estimates of the *m*^th^ mixture weight (*m* = 1,2,3) in cancer type *c*. Let θ^*^ represent the current value of all the remaining parameters, let *i* represent the sample number in cancer type *c* (1≤*i*≤*n*_c_) and let

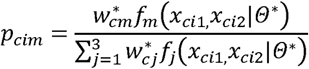

Then the EM updates are as follows:

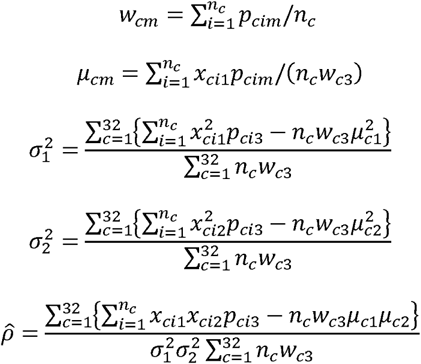

### Accounting for the uncertainty of the estimated pan-cancer correlation

Here *ρˆ* a pan-cancer measure of correlation between lncRNA and PC gene. For each correlation estimate, we calculate the standard error of the estimate (SE(*ρˆ*)) by bootstrapping with 100 bootstrap samples. This enables us to use a measure of the pan-cancer correlation that takes the uncertainty of the estimate into account, namely *ρˆ*/SE(*ρˆ*), which we refer to as the MCA score. Where lncRNA or PC gene expression signal is insufficient to calculate a correlation estimate, the cancer type is not considered further for this combination. In a significant number of cases, low expression of the lncRNA means the correlation cannot be estimated, and thus failure to calculate an MCA score for a specific PC gene. Where this occurs for over 50% of the PC genes, the lncRNA is not considered further, resulting in removal of a further 265 lncRNAs. Overall, 1833 lncRNAs have an MCA score for more than 50% of the 17088 PC genes.

### Weighted correlation network analysis (WGCNA)

To perform WGCNA [17], the R package “*WGCNA*” was applied as follows. First, a weighted lncRNA MCA score correlation network was constructed from the 1833 lncRNA by 17088 PC gene MCA score matrix using a soft thresholding power of 7 to which the MCA score correlation was raised to calculate adjacency. To aid choice of soft thresholding power we used the “pickSoftThreshold” WGCNA function with candidate powers 1-10, 12, 14, 16, 18 and 20. The power 7 was the lowest power for which the scale-free topology fit index reached 0.95 (Figure S3A, resulting in a network with mean connectivity of 5.94 (Figure S3B). Modules were then identified by average linkage hierarchical clustering of lncRNAs, and modules identified in the resulting dendrogram by the Dynamic Hybrid tree cut using signed topographical overlap matrix (TOM) and network types, a minimum module size of five, and a threshold for merging high correlated modules of 0.25. All other parameters were set to their default values.

### Signature enrichment analysis

Functional, cell type, transcription factor and disease type enrichment analyses were performed on each set of pro‐ and anti-module PC genes using Toppgene [48]. Significant enrichments were defined as those achieving False Discovery Rate less than 0.05 and signature overlap greater than two genes.

### Differential expression between tumour and normal samples

RNA-Seq raw FASTQ sequence files for TCGA matched normal samples across 24 cancer types were downloaded from CGHub, and gene expression estimates derived using the same procedure as for the tumour samples. 10 cancer types comprised of <10 samples after filtering so were removed from further analyses (Table S1). Differentially expressed lncRNAs (|log_2_FC|>1.0 and *p*<0.0001) between tumour and normal samples in each of the remaining 14 cancer types were detected using the Student’s *t*-test on FPKM expression estimates.

### De novo transcription factor motif discovery

Nucleotide sequences 1000bp upstream of each lncRNA were downloaded from Ensembl version 84 [49], and grouped according to module membership. Conserved motifs within these sequences from ME4 and ME5, and ME7-ME16 were then determined by a Weeder 2.0 [50] *de novo* search with default parameters. Modules without a coherent functional/cell type signature (ME1) or associated with transcriptional regulation only (ME2, ME3, ME6) were ignored. Motif matrices achieving scores >2.0 were then assessed for similarity with transcription factor binding sites contained within the JASPAR database using the JASPAR matrix alignment tool [23]. *De novo* matrices achieving >95% with a JASPAR matrix were deemed significant. Motifs associated with lncRNAs of ME13 were manually inspected using the Repeat Masker (http://www.repeatmasker.org) track on the University of California Santa Cruz (UCSC) Genome Browser [51].

### A novel approach to identify stromal cell specific lncRNAs

To further establish the stromal cell specificity of lncRNAs in ME12 and ME16, we used a novel approach to compare their expression in sample types that consist exclusively of tumour cells (stroma^low^) with fresh frozen TCGA patient samples that naturally contain a mixed population of tumour and stromal cells (stroma^high^). We reasoned that lncRNAs detected in stroma^high^ but not in stroma^low^ samples were likely stromal cell specific (for this purpose, immune cells are included in the definition of “stroma”). To represent stroma^low^ samples, we used 828 cell lines from the Cancer Cell Line Encyclopaedia (CCLE; Table S13) [52], and 57 PDX models [53], in which tumour had been separated from stroma using an *in silico* species-specific mapping strategy [53][54]. As expected, both stroma^low^ cohorts achieved a mean estimated tumour cell content of 99%±1%, compared to patient samples from TCGA where only 8/32 cancer types achieved median tumour cell content>90% (Table S12).

BAM files consisting of reads mapped to the human (GRCh37) genome were downloaded from the CGHub for the 828 cell lines representing 19 solid cancer types (Table S13). Only cancer types represented in the TCGA dataset were considered. FPKM values for each gene present in the human (GRCh38) GENCODEv19 annotation file were calculated as before using Cufflinks with library type defined as “fr-unstranded”.

RNA-Seq data for the 57 PDX models representing eight cancer types (25 lung, 12 breast, 7 colorectal, 3 endometrial, 6 ovarian, 2 pancreatic, 1 ampullary and 1 leukaemia) were downloaded from ArrayExpress (accession number: E-MTAB-3980), and tumour and stromal expression separated according to [53]. Note that the tumour components of 22/69 PDX models in the original dataset showed evidence of patient stroma retention (mRNA expression of CAF markers *FAP* or *CSPG4* log_2_ FPKM>2.0) so were ignored [53].

For the 1540 lncRNAs common to TCGA, CCLE and PDX datasets, we counted the number of tumour types in which the lncRNA was undetected in cell lines but detected in patient tumours (*x*), and the number of tumour types in which lncRNA was detected in patients regardless of cell line expression (*y*). Here, “detected” in patient tumours was defined as median FPKM>1.00 across the cancer type, and “undetected” in cell lines defined as median FPKM<0.50. 496 lncRNAs achieved *x*/*y* ≥ 0.50 and *x*>1, or *x*/*y*=1.00 and *x*=1, and therefore classed as undetected in cell lines and detected in patient tumours (set A). 768 lncRNAs were classed as undetected in our PDX cohort, achieving a median read count across the 57 models of zero (set B). 300 lncRNAs formed the union of sets A and B, and were therefore classed as stromal cell specific (SCS) achieving expression in patient tumours but low or undetectable expression in either cell lines or PDX models. SCS lncRNAs included *MEG3*, one of the few lncRNAs established as preferentially expressed in tumour stroma [55], thus adding confidence to our approach.

### esiRNA knockdown

esiRNAs were prepared as described in [56] using DEQOR [57] and primer3 [58] for optimized design of the template. An *in vitro* transcription kit (Thermo) was used to generate the dsRNA according to manufacturer’s instructions, followed by SureCut RNase III (NEB) digestion. After testing for complete digestion prior to use by agarose gel electrophoresis, esiRNAs were transfected into human primary fibroblasts at 5 ng per well in a total volume of 25μl. After 48 h, TGF‐ β1 (R and D Systems; 5 ng/ml) was added in serum-free medium. After a further 24hrs, fibroblasts were fixed in formaldehyde and monitored for αSMA induction using high content microscopy and using a FITC-conjugated anti‐ αSMA monoclonal antibody (Sigma)

## Acknowledgments

JRB is funded by Yorkshire Cancer Research (YCR SHEND01/02; http://yorkshirecancerresearch.org.uk). ICP is supported by a University of Malaya-Malaysian Ministry of Higher Education High Impact Research grant (UM.C/625/1/HIR/MOHE/DENT/22; https://www.mohe.gov.my/en). The results published here are in whole or part based upon data generated by The Cancer Genome Atlas managed by the NCI and NHGRI. Information about TCGA can be found at http://cancergenome.nih.gov. The funders had no role in study design, data collection and analysis, decision to publish, or preparation of the manuscript.

---

**Fig S1. Heatmap of eigen-lnc adjacencies.** Each row and column corresponds to one eigen-lnc. Within the heatmap, red indicates high adjacency (positive correlation) and green low adjacency (negative correlation) as shown by the colour legend.

**Fig S2. Typical three-component mixture distribution observed between PC and lncRNA gene expression.** The first two densities decay exponentially away from the ‐ and y-axes and the third distribution is a clearly separated bivariate Gaussian.

**Fig S3. Analysis of lncRNA-PC MCA score network topology for various soft-thresholding powers.** A. The scale-free fit index (y-axis) as a function of the soft-thresholding power (x-axis). B. mean connectivity (degree, yaxis) as a function of the soft-thresholding power (x-axis).

**Table S1. Number of TCGA patients contributing to this study across 32 cancer types.**

**Table S2: Module assignment and correlation of lncRNA association score profiles with the eigen-lncs.**

**Table S3: Eigen-lnc coefficients (PC-MA scores) contributed by each protein coding gene.**

**Table S4: ToppGene functional enrichment in pro-module protein coding genes.**

**Table S5: Module disease specificity.**

**Table S6: Evidence for FOS/JUN transcription factor binding sites in lncRNA promoters of module 7.** (a) Weeder motif scores. (b) Frequency matrix associated with top scoring motif (ATGAGTCATA). (c) Presence of top-scoring motif in ME7 lncRNAs. (d) Top 6 JASPAR database matches with top matrix hit (human-derived motifs only).

**Table S7: Enrichment of AP1 transcription factor binding sites in protein coding genes achieving PC-MA in module 7.**

**Table S8: Number and percentage lncRNAs in each module with ChipSeq evidence of SMAD3 occupancy.**

**Table S9: LncRNA detection in pre-clinical tumour models.** (a) Assessment of expression levels of each lncRNA in cell line and PDX tumour models. (b) Number and proportion of lncRNAs detected in cell lines/PDX models in each module.

**Table S10: Module-specific gene lists of extracellular-associated modules.** (a) ME16-specific. (b) ME12-specific.

**Table S11: Frequency of module-associated lncRNA dysregulation in cancer.** (a) LncRNAs differential expressed in each cancer. (b) LncRNAs differentially expressed in at least one cancer type and their dysregulation classification. (c) Number and proportion of each dysregulation class in each module.

**Table S12: PC-MA scores of genes in a reactive stroma signature.**

**Table S13: Number of CCLE cell lines contributing to this study across 19 cancer types.**

## References

1. Prensner JR, Chinnaiyan AM. The emergence of lncRNAs in cancer biology. Cancer Discovery 2011; 1: 391–407.

2. Gupta RA, Shah N, Wang KC, Kim J, Horlings HM, Wong DJ, et al. Long non-coding RNA HOTAIR reprograms chromatin state to promote cancer metastasis. Nature 2010; 464: 1071–1076.

3. Kogo R, Shimamura T, Mimori K, Kawahara K, Imoto S, Sudo T, et al. Long noncoding RNA HOTAIR regulates polycomb-dependent chromatin modification and is associated with poor prognosis in colorectal cancers. Cancer Research 2011; 71: 6320–6326.

4. Sørensen KP, Thomassen M, Tan Q, Bak M, Cold S, Burton M, et al. Long non-coding RNA HOTAIR is an independent prognostic marker of metastasis in estrogen receptor-positive primary breast cancer. Breast Cancer Res Treat; 2013, 142: 529–536.

5. Prensner JR, Iyer MK, Balbin OA, Dhanasekaran SM, Cao Q, Brenner JC, et al. Transcriptome sequencing across a prostate cancer cohort identifies PCAT-1, an unannotated lincRNA implicated in disease progression. Nat Biotechnology 2011; 29: 742–749.

6. Ji P, Diederichs S, Wang W, Boing S, Metzger R, Schneider PM, et al. MALAT-1, a novel noncoding RNA, and thymosin beta4 predict metastasis and survival in early-stage non-small cell lung cancer. Oncogene 2003; 22: 8031–8041.

7. Harrow J, Frankish A, Gonzalez JM, Tapanari E, Diekhans M, Kokocinski F, et al. GENCODE: the reference human genome annotation for The ENCODE Project. Genome Research 2012; 22: 1760–1774.

8. Iyer MK, Niknafs YS, Malik R, Singhal U, Sahu A, Hosono Y et al. The landscape of long noncoding RNAs in the human transcriptome. Nature Genetics 2015; 47: 199–208.

9. Yan X, Hu Z, Feng Y, Hu X, Yuan J, Zhao SD, et al. Comprehensive genomic characterization of long non-coding RNAs across human cancers. Cancer Cell 2015; 28: 529–540.

10. Li J, Han L, Roebuck P, Diao L, Liu L, Yuan Y, et al. TANRIC: An interactive open platform to explore the function of lncRNAs in cancer. Cancer Research 2015; 75: 3728–3737.

11. Guttman M, Amit I, Garber M, French C, Lin MF, Feldse F, et al. Chromatin signature reveals over a thousand highly conserved large non-coding RNAs in mammals. Nature 2009; 458: 223–227.

12. Cabanski CR, White NM, Dang HX, Silva-Fisher JM, Rauck CE, Cicka D, et al. Pan-cancer transcriptome analysis reveals long noncoding RNAs with conserved function. RNA Biol. 2015; 12: 628–642.

13. Liu Y, Zhao M. lnCaNet: pan-cancer co-expression network for human lncRNA and cancer genes. Bioinformatics 2016; 32:1595–1597.

14. Ashouri A, Sayin VI, Eynden JV, Singh SX, Papagiannakopoulos T, Larsson E. Pan-cancer transcriptomic analysis associates long non-coding RNAs with key mutational driver events. Nature Communications 2016; 13197.

15. Cogill SB, Wang L. Co-expression Network Analysis of Human lncRNAs and Cancer Genes. Cancer Informatics 2014; 13: 49–59.

16. Balbin OA, Malik R, Dhanasekaran SM, Prensner JR, Cao X, Wu Y-M, et al. The landscape of antisense gene expression in human cancers. Genome Research 2015; 25: 1068–1079.

17. Langfelder P, Horvath S. WGCNA: an R package for weighted correlation network analysis. BMC Bioinformatics 2008; 9: 559.

18. Vance KW, Ponting CP Transcriptional regulatory functions of nuclear long noncoding RNAs. Trends Genet. 2014; 30: 348–355.

19. Brunner AL, Beck AH, Edris B, Sweeney RT, Zhu SX, Li R, et al. Transcriptional profiling of long non-coding RNAs and novel transcribed regions across a diverse panel of archived human cancers. Genome Biology 2012; 13: R75.

20. Clark AT. The stem cell identity of testicular cancer. Stem Cell Rev. 2007; 3:49–59.

21. Loewer S, Cabili MN, Guttman M, Loh YH, Thomas K, Park IH, et al. Large intergenic non-coding RNA-RoR modulates reprogramming of human induced pluripotent stem cells. Nat Genet. 2010; 42: 1113–1117.

22. Mineo TC, Ambrogi V, Baldi A, Pompeo E, Mineo D. Recurrent intrathoracic thymomas: potential prognostic importance of cell-cycle protein expression. J Thorac Cardiovasc Surg. 2009; 138: 40–45.

23. Mathelier A, Zhao X, Zhang AW, Parcy F, Worsley-Hunt R, Arenillas DJ, et al. JASPAR 2014: an extensively expanded and updated open-access database of transcription factor binding profiles. Nucleic Acids Research 2014; 42 (Database issue): D142–147.

24. Zhang Y, He Q, Hu Z, Feng Y, Fan L, Tang Z, et al. Long noncoding RNA LINP1 regulates repair of DNA double-strand breaks in triple-negative breast cancer. Nature Structural & Molecular Biology 2016; 23: 522–530.

25. Zhang Y, Feng X-H, Derynck R. Smad3 and Smad4 cooperate with c-Jun/c-Fos to mediate TGF-beta-induced transcription. Nature 1998; 394: 909–913

26. Zhou C, York SR, Chen JY, Pondick JV, Motola DL, Chung DT et al. Long noncoding RNAs expressed in human hepatic stellate cells form networks with extracellular matrix proteins. Genome Medicine 2016; 8: 31.

27. Koinuma D, Tsutsumi S, Kamimura N, Taniguchi H, Miyazawa K, Sunamura M, et al. Chromatin immunoprecipitation on microarray analysis of Smad2/3 binding sites reveals roles of ETS1 and TFAP2A in transforming growth factor beta signaling. Mol Cell Biol. 2009; 29: 172–186.

28. Lamouille S, Xu J Derynck R. Molecular mechanisms of epithelial–mesenchymal transition. Nature Reviews Molecular Cell Biology 2014; 15: 178–196.

29. Verrecchia F, Vindevoghel L, Lechleider RJ, Uitto J, Roberts AB, Mauviel A. Smad3/AP-1 interactions control transcriptional responses to TGF-β in a promoter-specific manner. Oncogene 2001; 20: 3332–3340.

30. Davies M, Robinson M, Smith E, Huntley S, Prime S, Paterson I. Induction of an epithelial to mesenchymal transition in human immortal and malignant keratinocytes by TGF-beta1 involves MAPK, Smad and AP-1 signalling pathways. J Cell Biochem 2005; 95: 918–931.

31. Thuault S, Valcourt U, Petersen M, Manfioletti G, Heldin CH, Moustakas A. Transforming growth factor-beta employs HMGA2 to elicit epithelial-mesenchymal transition. J Cell Biol 2006; 174: 175–183.

32. Bakiri L, Macho-Maschler S, Custic I, Niemiec J, Guío-Carrión A, Hasenfuss SC, et al. Fra-1/AP-1 induces EMT in mammary epithelial cells by modulating Zeb1/2 and TGFβ expression. Cell Death Differ. 2015; 22: 336–350.

33. Yoshihara K, Shahmoradgoli M, Martínez E, Vegesna R, Kim H, Torres-Garcia W, et al. Inferring tumour purity and stromal and immune cell admixture from expression data. Nature Comms 2013; 4: 2612.

34. Subramanian A, Tamayo P, Mootha VK, Mukherjee S, Ebert BL, Gillette MA, et al. Gene set enrichment analysis, a knowledge-based approach for interpreting genome-wide expression profiles. PNAS 2005; 102: 15545–15550.

35. Zhou Y, Zhang X, Klibanski A. MEG3 noncoding RNA: a tumor suppressor. J Mol Endocrinol. 2012; 48: R45–53.

36. Mangone FRR, Brentani MM, Nonogaki S, Begnami MDFS, Campos AHJFM, Walder F, et al. Overexpression of Fos-related antigen-1 in head and neck squamous cell carcinoma. Int J Exp Pathol. 2005; 86: 205–212.

37. Farmer P, Bonnefoi H, Anderle P, Cameron D, Wirapati P, Becette V, et al. A stroma-related gene signature predicts resistance to neoadjuvant chemotherapy in breast cancer. Nature Medicine 2009; 15: 68–74.

38. Malanchi I, Santamaria-Martínez A, Susanto E, Peng H, Lehr H-A, Delaloye J-F et al. Interactions between cancer stem cells and their niche govern metastatic colonization. Nature 2012; 481: 85–89.

39. Liu J, Chen S, Wang W, Ning BF, Chen F, Shen W et al. Cancer-associated fibroblasts promote hepatocellular carcinoma metastasis through chemokine-activated hedgehog and TGF‐ pathways. Cancer Lett. 2016; 379: 49–59.

40. Franco OE, Shaw AK, Strand DW, Hayward SW. Cancer associated fibroblasts in cancer pathogenesis. Semin. Cell Dev. Biol 2010; 21: 33–39.

41. Kalluri R, Zeisberg M. Fibroblasts in cancer. Nat. Rev. Cancer 2006; 6: 392–401.

42. Jia CC, Wang TT, Liu W, Fu BS, Hua X, Wang GY, et al. Cancer-associated fibroblasts from hepatocellular carcinoma promote malignant cell proliferation by HGF secretion. PLoS ONE 2013; 8: e63243.

43. Mellone M, Hanley CJ, Thirdborough S, Mellows T, Garcia E, Woo J, et al. Induction of fibroblast senescence generates a non-fibrogenic myofibroblast phenotype that differentially impacts on cancer prognosis. Aging (Albany NY) 2016; 9: 114–132.

44. Bagordakis E, Sawazaki-Calone I, Macedo CC, Carnielli CM, de Oliveira CE, Rodrigues PC, et al. Secretome profiling of oral squamous cell carcinoma-associated fibroblasts reveals organization and disassembly of extracellular matrix and collagen metabolic process signatures. Tumour Biol. 2016; 37: 9045–9057.

45. Dobin A, Davis CA, Schlesinger F, Drenkow J, Zaleski C, Jha S, et al. STAR: ultrafast universal RNA-seq aligner. Bioinformatics 2013; 29: 15–21.

46. Trapnell C, Williams BA, Pertea G, Mortazavi A, Kwan G, van Baren MJ, et al. Transcript assembly and quantification by RNA-Seq reveals unannotated transcripts and isoform switching during cell differentiation. Nature Biotechnology 2010; 28: 511–515.

47. Johnson WE, Rabinovic A, Li C. Adjusting batch effects in microarray expression data using Empirical Bayes methods. Biostatistics 2007; 8: 118–127.

48. Chen J, Bardes EE, Aronow BJ, and Jegga AG. ToppGene Suite for gene list enrichment analysis and candidate gene prioritization. Nucleic Acids Res 2009; 37: W305–311.

49. Yates A, Akanni W, Amode MR, Barrell D, Billis K, Carvalho-Silva D, et al. Ensembl 2016. Nucleic Acids Res. 2016; 44: D710–D716.

50. Zambelli F, Pesole G, Pavesi G. Using Weeder, Pscan, and PscanChIP for the discovery of enriched transcription factor binding site motifs in nucleotide sequences. Curr. Protoc. Bioinform. 2014; 47: 2.11.1–2.11.31.

51. Kent WJ, Sugnet CW, Furey TS, Roskin KM, Pringle TH, Zahler AM, et al. The human genome browser at UCSC. Genome Res. 2002; 12: 996–1006.

52. Barretina J, Caponigro G, Stransky N, Venkatesan K, Margolin AA, Kim S, et al. The Cancer Cell Line Encyclopedia enables predictive modelling of anticancer drug sensitivity. Nature 2012; 483: 603–607.

53. Bradford JR, Wappett M, Beran G, Logie A, Delpuech O, Brown H, et al. Whole transcriptome profiling of patient derived xenograft models as a tool to identify both tumour and stromal specific biomarkers. Oncotarget 2016: 7: 20773–20787.

54. Bradford JR, Farren M, Powell SJ, Runswick S, Weston SL, Brown H, et al. RNA-Seq differentiates tumour and host mRNA expression changes induced by treatment of human tumour xenografts with the VEGFR tyrosine kinase inhibitor cediranib. PLOS ONE 2013; 8: 66003.

55. Zhang Z, Weaver DL, Olsen D, deKay J, Peng Z, Ashikaga T, et al. Long non-coding RNA chromogenic in situ hybridisation signal pattern correlation with breast tumour pathology. J. Clin. Pathol. 2016; 69: 76–81.

56. Theis M, Paszkowski-Rogacz M, Weisswange I, Chakraborty D, Buchholz F. (2015) Targeting human long noncoding transcripts by endoribonuclease-prepared siRNAs. J Biomol Screen. 2015; 20: 1018–1026.

57. Henschel A, Buchholz F, Habermann B. DEQOR: a web-based tool for the design and quality control of siRNAs. Nucleic acids research 2004; 32: W113–120.

58. Untergasser A, Cutcutache I, Koressaar T, Ye J, Faircloth BC, Remm M, et al. Primer3 - new capabilities and interfaces. Nucleic Acids Research 2012; 40: e115.

